# ASPIRE: the Amplicon Sequencing Profiler for Investigating Respiratory Ecosystems

**DOI:** 10.64898/2026.08.05.743000

**Authors:** Ryan J. McLaughlin, Steven Q. Chen, Anika Nag, Avery J. C. Noonan, Crista Bartolomeu, Scott A. Borden, Stephen Lam, Renelle Myers, Steven J. Hallam

## Abstract

**Summary:** Microbial communities inhabiting the respiratory tract contribute to health status through interactions with host physiology, immune function, and local environmental conditions. Advances in small subunit ribosomal RNA (SSU or 16S rRNA) gene amplicon sequencing enable culture-independent profiling of microbial communities as amplicon sequence variants (ASVs), revealing links between microbial dysbiosis and respiratory diseases, and the use of mass spectrometry to measure volatile organic compounds (VOCs) in exhaled breath shows emerging promise for biomarker discovery. Here we present ASPIRE, the Amplicon Sequencing Profiler for Investigating Respiratory Ecosystems, for processing, analyzing, and interpreting linked ASV-VOC data from respiratory microbiome studies and beyond. ASPIRE is designed to support scalable comparative analysis across respiratory sample types while preserving intermediate file outputs for inspection and reuse within a standardized file structure.

**Availability and implementation:** The complete workflow, test data, and required documentation are publicly available as a GitHub repository at https://github.com/hallamlab/ASPIRE. ASPIRE is implemented in Nextflow with a modular design architecture for reproducibility, extensibility and scalability.

## 1.0 Introduction

The human lung microbiome plays an important role in respiratory health and disease by influencing immune function and inflammatory responses (1). Advances in small subunit ribosomal RNA (SSU or 16S rRNA) gene amplicon sequencing have enabled culture-independent profiling of microbial communities as amplicon sequence variants (ASVs), revealing links between microbial dysbiosis and respiratory diseases such as COPD, asthma, and lung cancer (2-4). However, microbial composition alone does not directly capture metabolic activity in the airways. Breath biopsy, or breathomics, provides a non-invasive complementary approach by measuring a complex mixture of volatile organic compounds (VOCs) and other volatile molecules released into exhaled breath from various catabolic mechanisms. Disease-associated VOC signatures have emerged as promising biomarkers for respiratory diseases, including COPD and lung cancer, with the potential to facilitate diagnosis by reducing reliance on invasive tissue sampling (5). Emerging lines of evidence suggest potential interactions between the lung microbiome and volatilome (6, 7), and several cultivated respiratory tract pathogens including *Staphylococcus aureus* and *Pseudomonas aeruginosa* emit characteristic VOC profiles (8). Despite the potential of combined ASV and VOC data in monitoring respiratory health status, accessible workflows for integrated analysis remain limiting. The Amplicon Sequencing Profiler for Investigating Respiratory Ecosystems (ASPIRE) provides a solution to this problem. Implemented in Nextflow (9), ASPIRE accepts raw SSU rRNA gene amplicon sequencing reads and paired VOC measurements, performs ASV preprocessing, filtering, and taxonomic labelling along with correlation analysis to identify statistically significant features relevant to respiratory health status or other state changes associated with integrated ASV-VOC data sets.

## 2.0 Implementation

ASPIRE combines several open-source software applications, including cutadapt (v5.2)(10), fastp (v1.3.6)(11), seqkit (v2.13.0)(12), VSEARCH (v2.31.0)(13), BLAST (v2.17.0+)(14), SINA (v1.7.2)(15), QIIME2 (v2024.10.1)(16), decontam (v1.22.0)(17), ConQueR (v2.0) (18), vegan (v) MITOMASTER API (19), scikit-bio (v0.7.3)(20), scikit-learn (v1.9.1)(21), scipy (v1.17.1)(22) indicspecies (v1.8.0)(23), and SPIEC-EASI (1.1.2)(24), igraph (v2.3.4)(25), UMAP (v0.5.11)(26), HDBSCAN (v0.8.44)(27) together with custom Python3 (v3.11) and R (v4.5.3) scripts for data integration, visualization, and statistical analysis and is executed as a Nextflow DSL2 (v25.10.0)(9) workflow using paired-end SSU rRNA gene amplicon FASTQ files, a matched VOC abundance table, sample metadata, and a sequencing manifest (Figure 1A). Initially benchmarked using ASV data from the SPARK21 lung microbiome study (28) and paired VOC data from the SPARK21 integrated ASV-VOC study (29), ASPIRE also supports user-defined metadata fields for grouping and statistical analyses. Key aspects of the workflow are described below.

**Figure 1:**
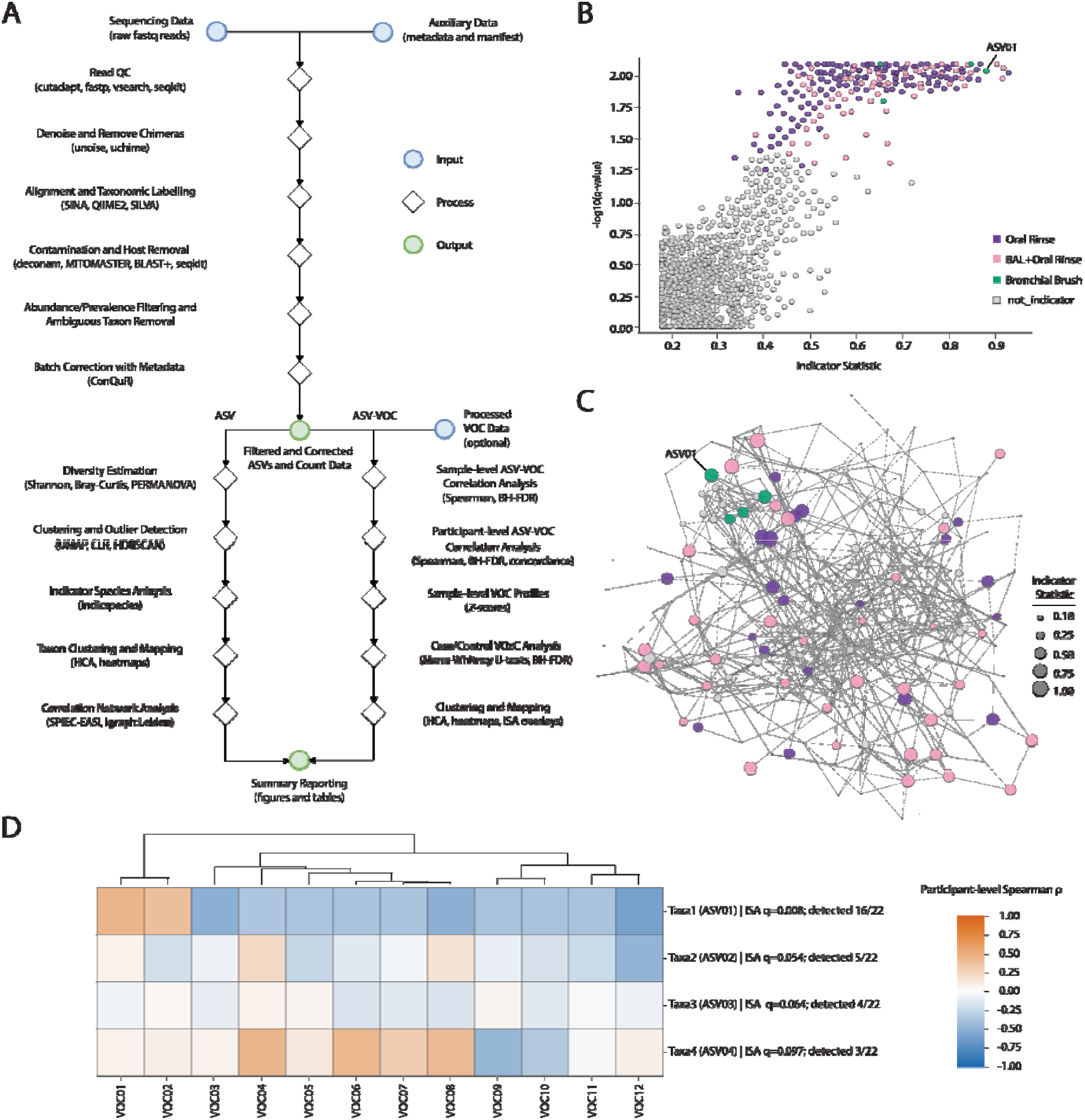
A) Schematic overview of the ASPIRE workflow, including ASV-only and ASV VOC branches. B) Multi-level indicator-species analysis (ISA): indicator statistic versus log10(q-value) illustrating ASVs significantly associated with respiratory sample types including oral rinse, bronchoalveolar lavage (BAL), and bronchial brushes (dashed line indicates FDR-adjusted q<0.05). C) SPIEC-EASI microbial co-occurrence network of ASVs, node color indicates inherited ISA status, and node size represents indicator statistic. D) ASV-VOC hierarchical cluster analysis (HCA) heatmap: normalized Spearman correlations for bronchial brush associated ASVs and VOCs, each ASV label contains its associated ISA q-value and detected prevalence in bronchial brush samples. ASV01 is labelled throughout subplots B-D to demonstrate explicit linkages between data products.

### 2.1 Read Quality Control and ASV Processing

Raw paired-end reads are primer- and quality-trimmed using Cutadapt (10) and fastp (11) then merged and quality-filtered using user-defined maximum expected error and sequence length thresholds with VSEARCH (13). Filtered reads are relabelled, concatenated, and dereplicated to construct the set of unique ASV sequences. ASVs are run through denoising and chimera removal with UNOISE (30) and UCHIME (31) functions implemented in VSEARCH. Quality-filtered reads are then mapped back to non-chimeric ASV centroids at 99.9% identity using VSEARCH to generate the initial ASV count matrix.

### 2.2 Taxonomic Annotation and Background Removal

ASVs are aligned to SILVA v138.2 (32) using SINA (15) to identify and filter for the target amplicon regions (ex. V4 region). ASV sequences are taxonomically classified using the QIIME2 (16) feature-classifier against the SILVA v138.2. Both SINA and QIIME2 formatted databased are downloaded automatically and stored for future use. Control-aware decontamination discriminates biological samples, technical negatives and biological/sample controls using the provided metadata table. Separate prevalence tests compare technical and biological controls with biological samples using the ‘decontam’ R package (17). The union of ASVs scoring below the configured threshold (default=0.1) in either enabled test are removed. Optionally, control subtraction is performed for remaining control background by subtracting the mean abundance of each ASV in user-defined control groups from each sample of interest. ASVs are then screened for human mitochondrial sequences and laboratory contaminants using the MITOMASTER API (19), BLAST+ (14), and custom Python scripts. The final microbial ASV table is filtered by sample representation, relative abundance, and taxonomy.

### 2.3 Metadata Integration and Quality Assessment

Metadata-linked ASV outputs are generated for downstream analyses and visualizations via customizable python scripts. Data-loss is accounted and visualized via a Sankey plot. ASV overlap across sample groups is visualized using Venn diagrams and UpSet plots. Optionally, batch correction, outlier analysis, and data transformations can be run at this stage. Batch correction, via the ‘ConQuR’ R package (18), produces a corrected count table which is then centered log-ratio (CLR) transformed and visualized with Uniform Manifold Approximation and Projection (UMAP) with Hierarchical Density-Based Spatial Clustering of Applications with Noise (HDBSCAN) diagnostics (26, 27). Outlier diagnostics combine Isolation Forest, one-class support-vector machines, and HDBSCAN with a configurable consensus vote (21).

### 2.4 Community Structure and Diversity Analysis

ASPIRE generates several different visualizations commonly used in microbiome studies, including relative abundance bubble plots, histograms, scatter plots representing UMAP embeddings, heatmaps and dendrograms. Collector’s curves, alpha diversity using the Shannon diversity index, and beta diversity using Bray-Curtis dissimilarity are calculated for microbial ASVs (20, 22). Participant-aware Shannon comparisons use paired Wilcoxon tests. Bray-Curtis beta diversity uses participant-blocked PERMANOVA with PERMDISP evaluated in parallel (9,999 permutations; seed 42).

### 2.5 Indicator Species Analysis and Microbial Network Analysis

Indicator species analysis (ISA) is performed using the R package indicspecies:multipatt (23) to identify taxa associated with sample type and disease status, retaining features with indicator statistic ≥0.25 and FDR-adjusted *q*<0.05 (Figure 1B). ASV and mitochondrial clustermaps are constructed across taxonomic ranks and incorporate indicator species annotations to identify sample-type-specific microbial signatures. Microbial co-occurrence networks are generated using SPIEC-EASI (24) with graphical lasso StARS model selection (Figure 1C). Additionally, igraphs(Leiden) (25) is run to compute ASV modularity within the network. Lastly, networks are overlayed with taxonomy, abundance, indicator species status, Leiden module assignment, and other metadata-defined groupings.

### 2.6 VOC-Branch ASV Filtering and ASV-VOC Integration

ASPIRE integrates matched ASVs, VOC measurements and metadata using sample and participant identifiers. ASVs are considered globally eligible when they reach ≥0.5% relative abundance in at least one sample, ≥5% prevalence and nonzero variance. The union of these ASVs and significant host-filtered sample-type ISA indicators defines the primary candidate set. Sample-level ASV-VOC associations are evaluated using Spearman rank correlations with two-sided p-values and Benjamini-Hochberg (BH) correction (22). For participant-level ASV-VOC inference, relative abundances and CLR-transformed abundances are calculated before within-participant averaging, with CLR evaluated as a sensitivity analysis. Pairwise Spearman correlations are assessed using two-sided participant-level permutation tests (9,999 permutations; seed 42), followed by BH correction within each normalization (22). Correlation direction and significance are compared between relative-abundance and CLR analyses to assess normalization concordance. Sample-level VOC profiles are standardized using per-VOC z-scores to describe relative elevations and depletions across samples. Case/control differences in participant-mean VOC abundance are evaluated using two-sided permutation Mann-Whitney U tests with BH correction and median differences as effect summaries (22). Clustering and visualization include hierarchical clustering analysis (HCA) of sample-level and participant-level ASV-VOC correlation matrices, with globally eligible, indicator-selected and user-defined grouping views (Figure 1D), alongside sample-level VOC z-score heatmaps. HCA uses average linkage and correlation distance, with Euclidean distance substituted when constant profiles occur (22). Case/control VOC differences are additionally visualized using differential-abundance bar plots with BH-adjusted significance annotations.

## 3.0 Conclusion

ASPIRE provides a scalable, flexible and reproducible workflow for integrated ASV-VOC analysis from respiratory microbiome studies that can be implemented on local, grid, or cloud-based infrastructure depending on data processing requirements. The workflow is designed to support comparative analysis across respiratory sample types providing a principled framework for linking microbial community structure with host-associated phenotypes. ASPIRE exposes core sequence processing, standard final-table preparation, and optional analyses through its YAML configuration. Read accounting and metadata summaries, Sankey plots, UpSet/Venn diagrams, collector curves, histograms, bubble plots, batch correction, outlier analysis, UMAP/HDBSCAN, dendrograms, heatmaps, clustermaps, indicator displays, network overlays, and integrated master tables summarize their respective inputs.

Initial benchmarking results using data from the SPARK21 lung cancer study indicate that ASPIRE can integrate microbiome profiles with VOC abundance data producing both global ASV-VOC association maps and focused indicator-species analyses in the context of respiratory health status. Although demonstrated here using a lung cancer cohort, the workflow is applicable to other respiratory conditions, including COPD and asthma, as well as environmental studies incorporating paired ASV-VOC data sets or other relevant metadata features. Future work should include more in-depth parameterization using larger, independently collected and appropriately controlled cohorts to corroborate potential associations with an eye toward biomarker discovery and validation.

## 4.0 Acknowledgments

This work was funded by the Terry Fox Research Institute and Canadian Cancer Society SPARK21 grant and provisioned with allocations from the Digital Research Alliance of Canada’s Arbutus Cloud resources and UBC’s Advanced Research Computing Sockeye cluster and Chinook cloud storage resources. A.J.C.N. was supported by the NSERC CREATE Ecosystem Services, Commercialization Platforms and Entrepreneurship (ECOSCOPE) training program at the University of British Columbia, as well as salary support from the Vancouver General Hospital Foundation.

