## Supplemental Information for "ASPIRE: the Amplicon Sequencing Profiler for Investigating Respiratory Ecosystems"

† Current affiliation and address

^*^ To whom correspondence should be addressed:

University of British Columbia, Department of Microbiology and Immunology

2552-2350 Health Sciences Mall, Vancouver, BC, Canada V6T, 1Z3

**Supplementary Material**

The ASPIRE workflow, quick-start guide, test data, and usage documentation are available at <https://github.com/hallamlab/ASPIRE>.
